# Disruption of *CYCLOPHILIN 38* function reveals a photosynthesis-dependent systemic signal controlling lateral root emergence

**DOI:** 10.1101/2020.03.11.985820

**Authors:** Lina Duan, Juan Manuel Pérez-Ruiz, Francisco Javier Cejudo, José R. Dinneny

**Affiliations:** Biology Department, Stanford University, Stanford, CA 94305; Department of Plant Biology, Carnegie Institution for Science, Stanford, CA 94305; Instituto de Bioquímica Vegetal y Fotosíntesis, Universidad de Sevilla and Consejo Superior de Investigaciones Científicas, Avda Américo Vespucio 49, 41092 Sevilla, Spain

## Abstract

Photosynthesis in leaves generates the fixed-carbon resources and essential metabolites that support sink tissues, such as roots [1]. One of these products, sucrose, is known to promote primary root growth, but it is not clear what other molecules may be involved and whether other stages of root system development are affected by photosynthate levels [2]. Through a mutant screen to identify pathways regulating root system architecture, we identified a mutation in the *CYCLOPHILIN 38* (*CYP38*) gene, which causes an accumulation of pre-emergent stage lateral roots, with a minor effect on primary root growth. CYP38 was previously reported to maintain the stability of Photosystem II (PSII) in chloroplasts [3]. *CYP38* expression is enriched in the shoot and grafting experiments show that the gene acts non-cell autonomously to promote lateral root emergence. Growth of wild-type plants under low light conditions phenocopied the *cyp38* lateral root emergence phenotype as did the inhibition of PSII-dependent electron transport or NADPH production. Importantly, the *cyp38* root phenotype is not rescued by exogenous sucrose, suggesting the involvement of another metabolite. Auxin (IAA) is an essential hormone promoting root growth and its biosynthesis from tryptophan is dependent on reductant generated during photosynthesis [4,5]. Both WT seedlings grown under low light and *cyp38* mutants have highly diminished levels of IAA in root tissues. The *cyp38* lateral root defect is rescued by IAA treatment, revealing that photosynthesis promotes lateral root emergence partly through IAA biosynthesis. Metabolomic profiling shows that the accumulation of several defense-related metabolites are also photosynthesis-dependent, suggesting that the regulation of a number of energy-intensive pathways are down-regulated when light becomes limiting.

## Results and Discussion

### *CYP38* is necessary for lateral root emergence

Lateral roots form through oriented cell divisions within the pericycle tissue layer, hidden 3-layers deep inside the parent root [6]. The process of lateral root primordia emergence through these outer tissues involves communication between the organ primordia and overlying cell layers and is mediated by the hormone auxin [7,8]. To identify factors that play a role in this process, we conducted a mutant screen and characterized a recessive mutant (originally named *prematurely stunted lateral organs, presto*) that exhibits a severe reduction in lateral root density under standard growth conditions, while primary root growth is also significantly reduced, but to a lesser extent (Figure. 1A-C). Next-generation EMS mutation mapping using sequencing was used to identify the causative genetic locus, *CYPLOPHILLIN 38* (*CYP38*) (Figure. S1B). A T-DNA allele (*cyp38-5)* with an insertion in the sixth exon phenocopied the EMS allele (*cyp38-4*) (Figure. 1A-C), which has an early stop codon in the second exon (Figure. 1D). Expression of *CYP38* using its native promoter fully complement *cyp38-4* (Figure. 1E), indicating that *CYP38* is indeed the causative gene.

**Figure 1.**
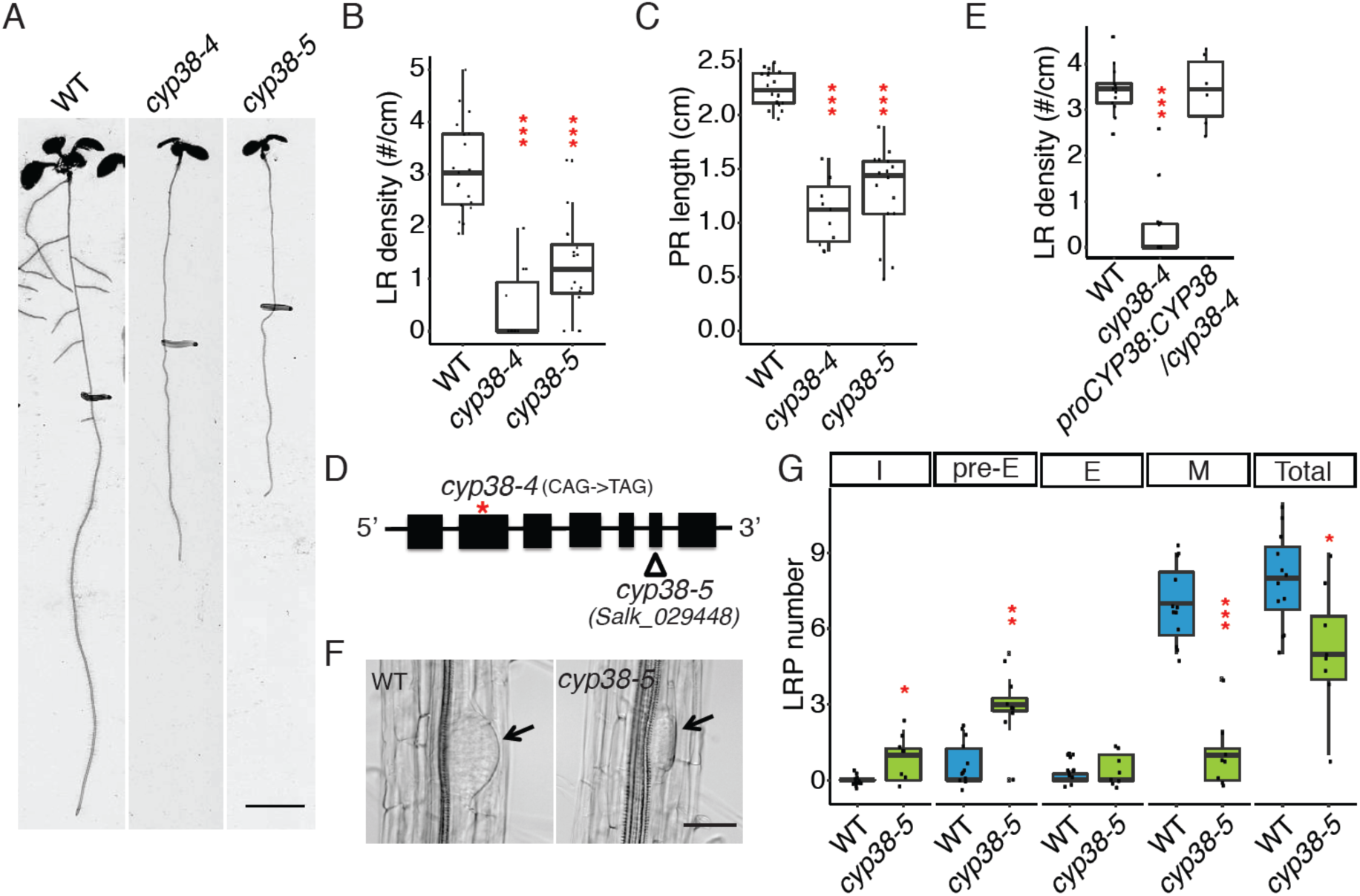
CYP38 is required for lateral root emergence. (A) Scanned root image of 11 dpg WT-0, *cyp38-4* and *cyp38-5*. Scale bar = 5 mm. (B, C) Quantifications of lateral root density (B) and primary root length (C) in two alleles of *cyp38*. (∗∗∗*P* < 0.001, pairwise t-test with Hochberg correction, n>10) (D) Diagram describing the mutation position of *cyp38-4* and T-DNA insertion position of *cyp38-5*. (E) Quantifications of LR density of the EMS allele, *cyp38-4*, and the complemented line. (∗∗∗*P* < 0.001, pairwise T-test with Hochberg correction, 6 individual complemented T1 plants are used). (F) Microscopic images of cleared WT and *cyp38-5* roots. Arrowheads point to pre-emerged lateral root primordia. Scale bar = 50 μm. (G) Quantifications of different stages of lateral root primordia in WT and *cyp38-5*. I represent the initiation stage, pre-E represents pre-emergence stage, E represents emerged lateral root stage, M represents matured lateral roots, and Total represents the sum of all 4 stages. (∗*P* < 0.05, ∗∗*P* < 0.01, ∗∗∗*P* < 0.001, pairwise T-test with Hochberg correction, n>8)

To determine the developmental basis for the *cyp38* phenotype, we classified lateral root growth into 4 distinct stages: initiation (I), pre-emergence (Pre-E), emergence (E), and maturation (M) (Figure. S1A). Compared to wild type (WT), where the majority of lateral roots have typically progressed to the maturation stage in 8-dpg (days post germination) seedlings, we found that the *cyp38-5* mutants had a significant enrichment of pre-emergence stage lateral roots (Figure. 1G). Lateral root primordia exhibited a flattened-shape in *cyp38-5* instead of the typical dome shape observed in WT (Figure. 1F). This defect has also been observed in other mutants that specifically affect the emergence stage [9]. It is likely that initiation and/or founder cell specification stages are also disrupted to account for the full reduction in lateral root development.

### CYP38 is localized to the chloroplast and regulates lateral root development through shoot-to-root signaling

As previously reported, *CYP38* encodes a cyclophilin-related protein important in the assembly and stability of PSII under high light [3,10,11]. However, limited characterization of the tissue-specific and subcellular localization of the protein has been performed. A transcriptional fusion, *proCYP38::erGFP*, and quantitative RT-PCR indicated that the gene is expressed predominantly in shoot tissues (mesophyll and hypocotyl) (Figure. 2A, Figure. S2A). Transient expression of a Ypet-tagged translational fusion to *CYP38* in Arabidopsis protoplasts showed a chloroplast-specific localization pattern (Figure. 2B). In stably transformed plants, only chloroplast or plastid localization was observed (Figure S2B). These data are consistent with a plastid-specific function for CYP38 in Arabidopsis.

**Figure 2.**
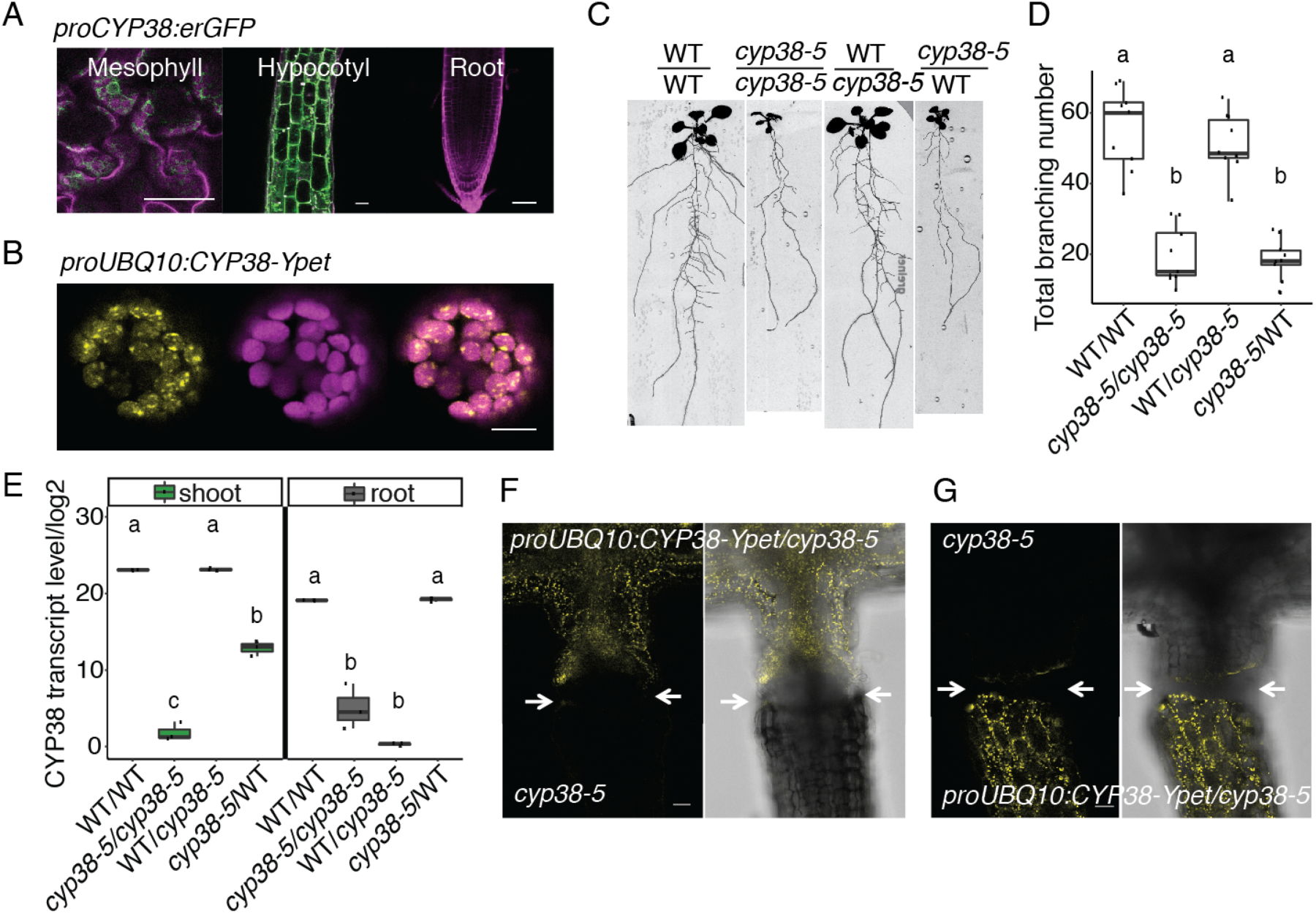
CYP38 is localized in chloroplast and regulates root branching through systemic signals. (A) Confocal images of *proCYP38:erGFP* transgenic line. Magenta indicates autofluorescence of chloroplasts in mesophyll cells together with propidium iodide (PI) staining in hypocotyl and root, while green indicates Green Fluorescent Protein (GFP). Scale bars = 50 μm. (B) Confocal images of WT protoplast that was transiently transformed with *proUBQ10:CYP38-Ypet*. Magenta indicates chloroplast autofluorescence, while yellow indicates Ypet fluorescence. Scale bars = 10 μm. (C) Scanned root images of the 17-day-old grafted plants. (D) Quantification of total branch number in 4 different grafted combinations. Significantly different groups are indicated with letters (*P* < 0.05), n>8. (E) Quantitative RT-PCR for *CYP38* transcript level in both shoot and root tissues of the grafted plants. The numbers are presented as log2 transformed values. Significantly different groups are denoted with letters (*P* < 0.05) and analyzed separately for shoot and root samples, n=3. (F, G) Confocal images of the hypocotyl junction in grafted plants. *proUBQ10:CYP38-Ypet* is used as the scion in (F), or used as the rootstock in (G). Arrowheads mark the grafting junction. Scale bars = 50 μm.

We next used grafting to resolve whether *CYP38* function in the shoot or root is responsible for the lateral root growth defects observed. Both wild type and *cyp38-5* mutant scions were grafted to WT or *cyp38-5* root stocks. Root branching was quantified 12 days post-grafting. Because we observed senescence of the primary root tip after formation of the graft junction, we quantified the total number of branches formed instead of lateral root density. Interestingly, grafting of a WT scion on to a *cyp38-5* root stock significantly rescued the lateral root emergence defect while a *cyp38-5* scion caused the WT root stock to exhibit a *cyp38*-like phenotype (Figure. 2C, D). These data suggest that the lateral root emergence phenotype is determined primarily by the genotype of the shoot.

As previously reported in tomato, the CYCLOPHILIN protein DIAGETROPICA promotes root branching by moving from the shoot to root [12,13]. To investigate whether *CYP38* mRNA or protein is able to translocate across the shoot-root junction, we compared expression in shoot and root tissues of the 4 grafted combinations described above. Interestingly, the *CYP38* transcript remained low in the WT/*cyp38-5* rootstock despite lateral root development being rescued. The transcript level in *cyp38-5*/WT scion is increased relative to the *cyp38-5* mutant root system, however, the levels are very low compared with the WT shoot (Figure. 2E). Thus, we conclude that there is little to no movement of *CYP38* mRNA across the shoot-root junction. Similarly, we also examined protein movement by grafting the shoot of *proUBQ10:CYP38-Ypet/cyp38-5* transgenic lines with a *cyp38-5* mutant root or shoot. Confocal microscopy was used to detect yellow fluorescence at the shoot-root junction after the graft junction formed successfully (about 5∼7 days post-grafting). No significant fluorescence could be detected across the grafted junction (Figure. 2F, G). In summary, the chloroplast localization pattern of CYP38 is consistent with previous findings on the importance of CYP38 in photosystem stability. The limited mobility of both mRNA and protein indicates that CYP38 promotes lateral root emergence non-autonomously from photosynthetic shoot tissues.

### Photosystem stability and chloroplast redox state regulate lateral root emergence

To determine whether the defect in lateral root emergence in *cyp38* mutants is a result of changes in overall photosynthetic activity, we firstly examined what effect light intensity had on root architecture. We manipulate light intensities layers of window screen, which we found reduces the intensity of the light without affecting the spectrum, particularly the red/far red light ratio (Figure. S3A, B). Arabidopsis seedlings were grown under standard light conditions (∼140 μE m^−2^ s^−1^) for 5 days and then transferred to several reduced light intensities for another 4 days. A severe reduction in visible lateral roots was observed when light intensity declined (Figure. 3A, B), while primary root growth was only moderately affected (Figure. 3C). We found that as the light intensity declined, lateral roots were increasingly enriched at the pre-emergence stage (Figure. 3D), indicating that this response reflected a specific inhibitory effect on the process of lateral root emergence rather than a general reduction in growth. Lateral root development is not necessarily always most restricted when seedlings growth is inhibited as our previous study showed that treatment with 1-aminocyclopropane-1-carboxylic acid (ACC) inhibited primary root growth more effectively than lateral roots [14].

**Figure 3.**
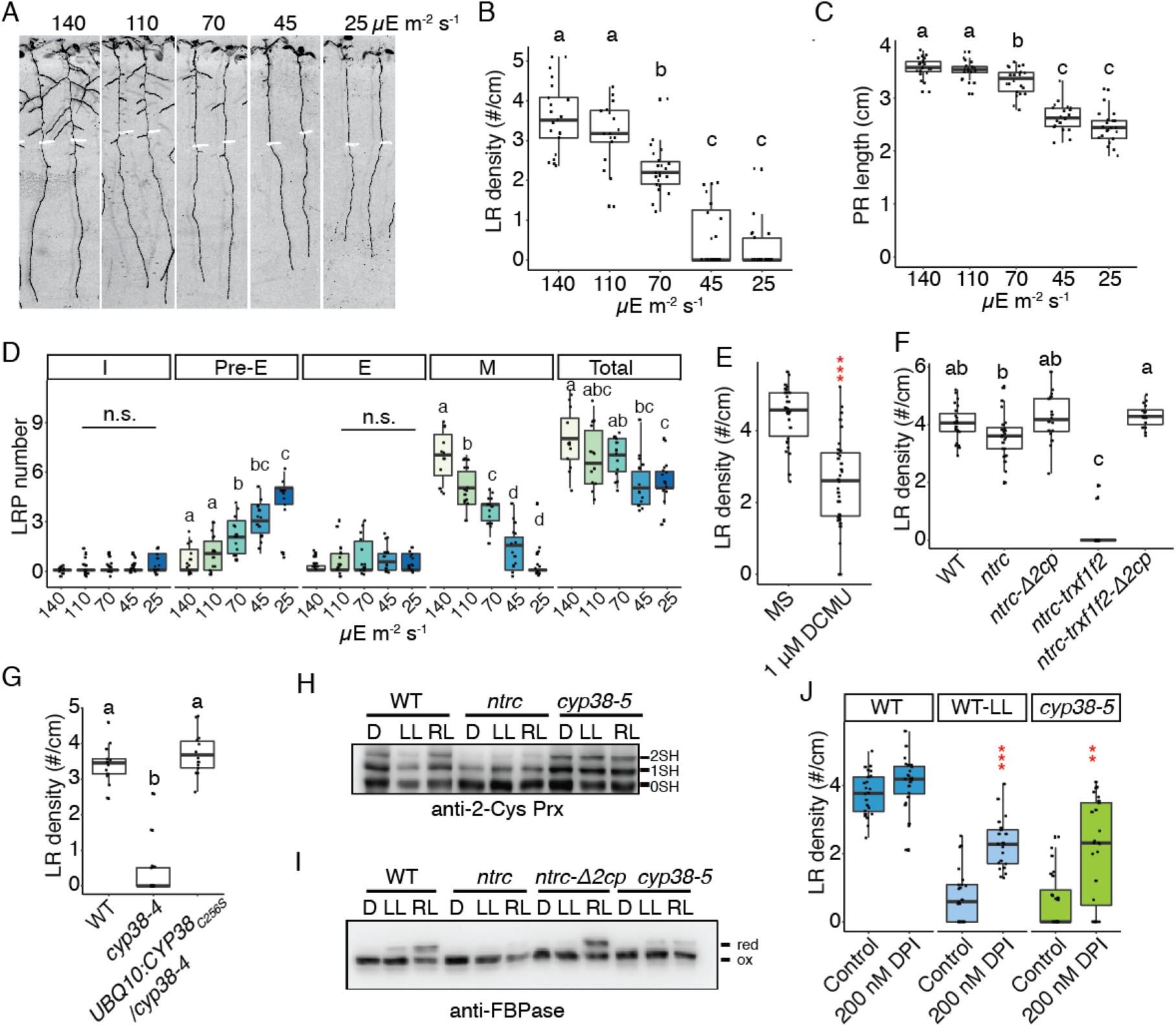
Chloroplast redox status regulates lateral root emergence. (A) Scanned root image of 9 dpg WT seedlings grown under 140 μE light for 5 days and then transferred to different light intensities for another 4 days. (B, C) Quantification of lateral root density (B) and primary root length (C) in WT under different light intensities. Significantly different groups are indicated with letters (*P* < 0.05), n>16. (D) Quantification of different stages of lateral root primordia in WT under different light intensities. I represent the initiation stage, pre-E represents pre-emergence stage, E represents emerged lateral root stage, M represents matured lateral roots, and Total represents the sum of all 4 stages. Significantly different groups are indicated with letters (*P* < 0.05), n>12. (E) Quantification of lateral root density in WT with 1 μM DCMU treatment. (∗∗∗*P* < 0.001, pairwise T-test, n>30) (F) Quantifications of lateral root density in *ntrc, ntrc-trxf1f2, ntrc-trxf1f1-*Δ*2cp* mutants under standard growth conditions. Significantly different groups are indicated with letters (*P* < 0.05), n>16. (G) Quantification of lateral root density in WT, *cyp38-4*, and complemented lines with cysteine to serine mutation in *CYP38* coding sequence. Significantly different groups are indicated with letters (*P* < 0.05), n>13 individual T1 complemented lines. (H, I) Wild type and mutant plants, as indicated, were grown under long-day conditions for 4 weeks at a light intensity of 125 μE m^−2^ s^−1^. *In vivo* redox status of 2-Cys Prx (H) and FBPase (I) proteins was determined at the end of the dark period (D) and after 30 min of illumination at 50 μE m^−2^ s^−1^ (LL) or 175 μE m^−2^ s^−1^ (RL). 0SH, 1SH and 2SH indicate reduction of zero, one or the two cysteine residues, respectively, of 2-Cys Prx (H). Red indicates reduced state and ox marks oxidized state of FBPase (I). (J) Quantifications of lateral root density in WT, *cyp38-5* under standard light conditions (140 μE m^−2^ s^−1^) and WT under low light (25 μE m^−2^ s^−1^), with or without 200 nM DPI treatment. (∗∗*P* < 0.01, ∗∗∗*P* < 0.001, two-way ANOVA comparing WT under control and DPI treatment, n>21).

To investigate whether the specific developmental response to light intensity is due to the direct perception of light by the root system in our tissue culture conditions, we utilized the GLO-Roots system to recapitulate a more physiologically relevant condition where roots are shielded from light and grown in soil [15]. Using this system, we observed a significant reduction in the number of lateral roots in *cyp38-5* mutant plants compared to wild type; a similar effect was observed in plants grown under low-light conditions (Figure. S3C, D). These data indicate that in both tissue culture conditions and soil conditions, the amount of light perceived by the shoot has a specific promoting effect on lateral root development.

To directly test whether the activity of the photosynthetic apparatus is required for lateral root emergence, we treated seedlings with 3-(3,4-dichlorophenyl)-1,1-dimethylurea (DCMU), which is a well-characterized inhibitor of electron transport between PSII and PSI. We observed significant inhibition in lateral root density (Figure. 3E), similar to the effect of low light (Figure. 3B).

We hypothesized that the products of the electron transport chain, such as NADPH or other downstream reducing agents may play important roles in modulating lateral root development. Consistent with this hypothesis, a reduction in lateral root growth was reported in mutant plants devoid of NADPH-dependent thioredoxin (Trx) reductase C (NTRC) [16], a chloroplast enzyme [17] that plays a central role in redox regulation of chloroplast proteins. NTRC is the most efficient electron donor to 2-Cys peroxiredoxins (Prxs), thereby the NTRC-2-Cys Prx redox system maintains the reductive capacity of chloroplast Trxs allowing the proper regulation of their targets [18]. To further characterize the contribution of these redox systems, NTRC-2-Cys Prx and Trxs, to lateral root development, we evaluated the various mutants reported, which have impaired redox homeostasis to different extents [18]. Interestingly, although only mild reduction in lateral root emergence was observed in *ntrc*, a strong inhibition was observed in *ntrc-trxf1f2* triple mutants (Figure 3F). This strong defect was completely rescued by the *ntrc-*Δ*2cp* and *ntrc-trxf1f2-*Δ*2cp* mutants, in line with previous results showing that decreased levels of 2-Cys Prxs restore chloroplast redox homeostasis of NTRC-deficient mutants [18].

Chloroplast cyclophilins have been identified as binding partners of Trx [19]. As reported by Vasudevan et al. [10], the mature CYP38 protein contains a single cysteine at position 256 in the cyclophilin domain. We checked whether this cysteine could mediate protein dimerization, through a disulfide bridge, by visualizing the protein on a non-reducing SDS-PAGE gel, however, no apparent dimerization band was found (Figure. S3E). We also mutated this cysteine to serine and transformed the mutated coding sequence back into *cyp38-4*. Full rescue of the mutant phenotype was observed (Figure. 3G), indicating that the function of CYP38 is not dependent on the cysteine and the protein is unlikely to be redox regulated. Because CYP38 has been identified as a potential partner of 2-Cys Prxs [19], we determined whether this cyclophilin acts as an upstream regulator of these peroxidases. Thus, we evaluated the redox status of 2-Cys Prxs and fructose bisphosphatase (FBPase), included here as a well-established target of the Trx pathway, in mutant plants lacking CYP38. While no difference was observed for 2-Cys Prxs (Figure 3H), light-dependent FBPase reduction was similarly impaired in *cyp38* and *ntrc* mutants under regular light intensity (Figure 3I). These data indicate that the redox status of some chloroplast proteins may be dependent on CYP38 activity and these proteins may overlap with Trx targets.

We next evaluated photosynthetic performance of CYP38-deficient mutants. Similar to the *ntrc* mutant, here included for comparison, quantum yield of photosystem II (Y(II)) was diminished in *ntrc* and *cyp38* plants (Figure S3F), indicating that both mutants had similarly affected the efficiency of light energy utilization. Interestingly, energy dissipation through non photochemical quenching (Y(NPQ)) was higher in *ntrc* than in the *cyp38* mutants (Figure S3G), whereas a significant elevation of Y(NO), which is an indicator of non-regulated energy dissipation, was only observed in *cyp38* (Figure S3H). Overall, these photosynthetic parameters suggest that the *cyp38* mutant does not only have reduced energy conversion efficiency, but is also inefficient in photodamage protection through heat dissipation. These data indicate potential side reactions such as excess ROS production in *cyp38*, which coincides with previous findings that higher H_2_O_2_ levels occur in this mutant [20]. Consistent with this hypothesis, adding diphenyleneiodonium chloride (DPI), which inhibits ROS production, partially rescues the lateral root defect in *cyp38* under standard light conditions, as well as WT root growth under low light conditions (Figure. 3J). ROS levels in lateral root primordia are important in regulating cell proliferation and differentiation in the primary root, and also facilitate the establishment of a dome shaped primordia [9]. Together, our observation suggests that the perturbed balance in chloroplast redox status may non-autonomously limit lateral root emergence.

### Sucrose and *HY5*-dependent signaling pathways regulate lateral root emergence independently from *CYP38*

Photosynthesis-derived sucrose serves as a long distance signal to control root growth [2]. Lateral root development can also be promoted by aerial uptake of sucrose from the media [21]. We tested whether sucrose is limiting for lateral root growth in *cyp38* mutants by supplementing our media with 1%, 2% and 3% sucrose. Importantly, the addition of sucrose had little effect on lateral root density (Figure S4A). Sucrose did not significantly rescue the inhibitory effects of low light either (Figure S4B).

*ELONGATED HYPOCOTYL 5* (*HY5*) acts as a mobile signal mediating root-to-shoot communication in response to light quantity and quality [22–24]. *hy5* mutants exhibit root systems with reduced primary root length, however, lateral root development is accelerated, relative to WT [23]. We found that *hy5* mutants showed similar lateral root density to WT under standard light conditions, while they showed a partial rescue of lateral root growth under low light conditions (Figure S4C, D). These data suggest that *HY5* mediates light-dependent growth-inhibition of lateral root development, which is similar to findings of a recent study [23]. To determine whether *HY5* acts in the same genetic pathway as *CYP38*, we compared the phenotypes of *hy5* and *cyp38* single and double mutants. While under standard light conditions the *hy5 cyp38* double mutant had an indistinguishable phenotype from the *cyp38* and *hy5* single mutants, under low light conditions the double mutant showed an intermediate reduction in lateral root development between that of *hy5* and *cyp38*, which we interpret as an additive effect (Figure S4C, D). Together these data suggest that *CYP38* and *HY5* likely regulate root growth through independent pathways. This is consistent with work showing that *HY5* likely mediates its effect on lateral root growth directly through local induction of *HY5* expression in light-exposed root tissues [23].

### Downregulation of auxin biosynthesis and signaling mediate the *cyp38* phenotype in roots

It is well known that auxin is a necessary signaling molecule throughout lateral root development [8,25,26]. To understand how the lateral root defect in *cyp38* mutants may be mediated by altered activity of the auxin pathway, we firstly measured IAA levels in the mutant and WT under varying light intensity. IAA accumulates to a higher level in the root than shoot in 10-day-old seedlings, and its abundance in root tissues is significantly reduced in the *cyp38-5* mutant as well as under low light conditions (Figure 4A). Using grafted plants, we found that IAA levels were highest in roots when the genotype of the shoot was WT, compared to plants where the shoot was grafted from *cyp38-5* mutants (Figure 4B). These data suggest that auxin biosynthesis in the shoot, and transmission to the root, is promoted by light intensity and *CYP38* function. An alternative hypothesis that we can not exclude is that an unknown signal mediates shoot to root communication and induces auxin biosynthesis locally in the root.

**Figure 4.**
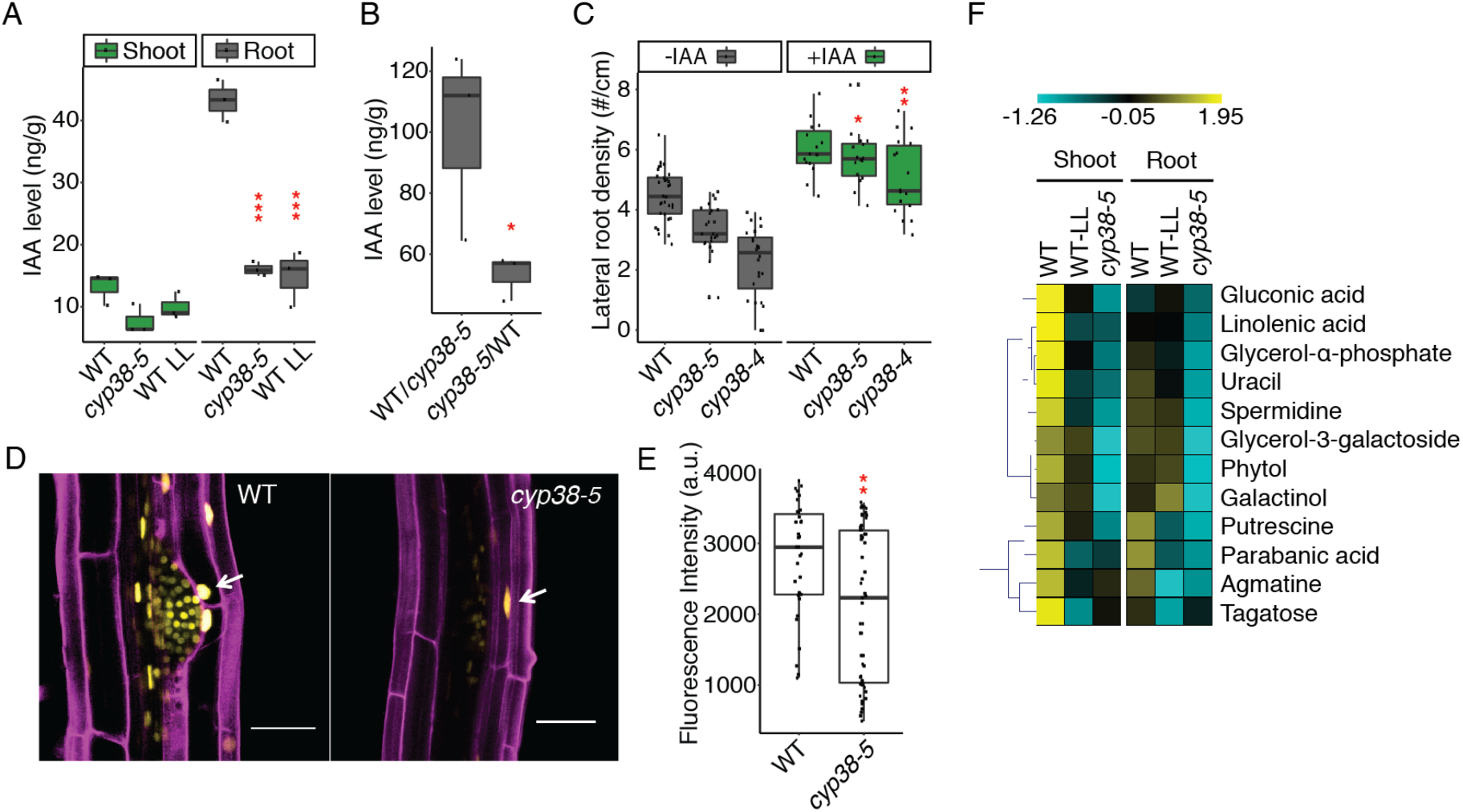
Auxin level and signaling and defense related metabolites are affected in cyp38. (A) Measurements of IAA levels in WT and *cyp38-5* under standard light and WT under low light in shoot and root tissues. (∗∗∗*P* < 0.001, pairwise T-test with Hochberg correction, n=3) (B) Measurements of IAA level in the rootstock of WT/*cyp38-5* and *cyp38-5*/WT grafted plants. (∗*P* < 0.05, pairwise T-test with Hochberg correction, n=3) (C)Quantification of lateral root density in WT and *cyp38* mutant with or without 0.5 μM IAA treatment. (∗*P* < 0.05, ∗∗*P* < 0.01, two-way ANOVA comparing with WT under control OR IAA treatment, n>15) (D) Confocal images of *pDR5rev::3XVenus-N7* in lateral root primordia in WT and *cyp38-5*. Arrowheads point to the expression in the cortex tissue layer surrounding a lateral root primordium. Scale bars = 50 μm. (E) Measurements of the fluorescence intensity of *pDR5rev::3XVenus-N7* in cortex cells surrounding lateral root primordia in WT and *cyp38-5*. (∗∗*P* < 0.01, pairwise T-test with Hochberg correction, n>36) (F) Heatmap of the subset of metabolites that show significant decrease in *cyp38-5* as well as IN WT under low light (one-way anova with *P* < 0.01, n=3).

Using a transcriptional reporter of auxin response, *pDR5rev::3XVENUS-N7 [27]*, we found that reporter activity in the primary root tip is similar between WT and *cyp38* while it is dramatically reduced in more distal mature tissues of *cyp38* (Figure. S5A, B). Auxin accumulates in the cortex cells surrounding lateral root primordia and induces expression of cell wall degrading proteins that facilitate the emergence of lateral root primordia [7]. We quantified *pDR5rev:3XVENUS-N7* expression in these cortical cells, and observed a significant reduction (Figure. 4D, E). Consistent with these observations, exogenous IAA treatment largely rescued lateral root emergence in *cyp38* (Figure. 4C). The conversion of Indole Pyruvic Acid (IPA) into IAA is dependent on NADPH-dependent reduction [28] and it may be that reduced photosynthetic output limits auxin biosynthesis.

### Defense-related metabolites are modulated by photosynthetic activity

Metabolic processes such as carbon fixation, starch metabolism and amino acid synthesis are affected by chloroplast redox status [29,30]. To identify metabolites that may change in abundance as a consequence of changes in light intensity or *CYP38* activity, we examined the levels of primary metabolites in shoot and root tissues of WT seedlings grown under standard or low-light conditions, as well as *cyp38* mutants grown under standard conditions (Figure. S5C, Table S1). Surprisingly, a significant number of the metabolites with reduced levels under low light conditions or in *cyp38* mutants are known biotic stress cues (Figure. 4F, Table S2): gluconic acid, which is derived from glucose, is associated with antifungal compound production [31]; glycerol-3-phosphate, which acts as a mobile factor for inducing systemic immunity in plants [32]; polyamines, including putrescine and spermidine, which regulate growth and also have important roles in modulating biotic and abiotic stress [33,34]; and linolenic acid, which is a precursor of the defense signaling molecules jasmonic acid and OPDA. Linolenic acid promotes lateral root development [35]. Under our conditions we found that exogenous application of linolenic acid to seedlings had no significant effect on WT under standard conditions, but partially rescued lateral root emergence in *cyp38* (Figure. S5D), suggesting it may also mediate changes in root architecture downstream of *CYP38*.

Together, our results reveal the interaction between photosynthesis and energy-intensive biological programs such as growth and defense. While many pathways are downregulated when photosynthetic activity of the shoot is lowered, the response of systemic organ systems is highly specific, with changes in root architecture that favor deeper root systems with fewer branches and a diminished defense response. These responses likely allow the plant to conserve fixed carbon resources to survive when total photosynthetic output becomes limiting.

## Methods details

### Plant materials

The *cyp38-4* mutant allele was derived from an EMS mutagenized population of Arabidopsis thaliana, Col-0. The *cyp38-5* allele was obtained from ABRC with the stock number SALK_029448. The *hy5* mutant was obtained from ABRC with the stock number SALK_096651. The *ntrc, ntrc-*Δ*2cp, ntrc-trxf1f2*, and *ntrc-trxf1f2-*Δ*2cp* lines are previously described [18]. Auxin reporter, *pDR5::3XVENUS-N7* was previously described [27]. Arabidopsis thaliana ecotype Columbia (Col-0) was used as the reference strain throughout the entire study.

### Plant growth conditions

Sterilized seeds were grown on sterile 0.7% Gelzan media containing 1X Murashige and Skoog nutrients (MSP01-50LT, Caisson) and 1% sucrose. Seedlings were grown under standard light intensity (140 μE m^−2^ s^−1^) for 5-6 days before transfer to standard media supplemented with chemicals, or reduced light conditions for 3-5 days. The position of the root tip was marked at the time of transfer. Supplements include IAA, DCMU, DPI and Linolenic acid (Sigma-Aldrich). Growth of seedlings was performed in a Percival CU41L4 incubator at a constant temperature of 22°C with long-day lighting conditions (16 hours light and 8 hours dark). Plates were sealed with both parafilm and micropore tape (3M) as previously described [14].

### Transgene construction

Golden gate cloning was used to assemble various DNA fragments into pSE7 vector [36]. The *proCYP38:CYP38* construct was generated by cloning a 5.1 kb genomic CYP38 fragment into Zero Blunt TOPO vector with the primers listed in Table S3. A 3 kb promoter region was used to generate the *proCYP38:erGFP* reporter construct. A VENUS derived fluorescence protein, Ypet [37], 2.5 kb *UBQ10* promoter and *CYP38* coding sequence was used to assemble *proUBQ10:CYP38-Ypet*. Site-directed mutagenesis was used to introduce Cys to Ser mutation in *CYP38* coding sequence in order to generate *proUBQ10:CYP38_c256s*. All primers used for cloning are listed in Table S3. Transgenic plants were generated by a standard floral dip method into the *cyp38-4* background. Seeds were harvested from transformed plants and selected visually based on mCherry fluorescence using an M165 FC fluorescence microscope (Leica).

### Microscopic analysis

For quantitation of lateral root developmental stages, roots were mounted in a modified Hoyer solution (chloral hydrate:water:glycerol in proportions 8:2:1, g/ml/ml), then imaged using a Leica DMI6000 inverted compound microscope. For confocal microscopy, roots were mounted in 5 μg/ml propidium iodide (PI) solution (Thermal Fisher), and imaged using a Leica SP8 point-scanning confocal microscope. The 488 nm laser was used to detect GFP and PI staining, and 512 nm laser was used for Ypet fluorescence excitation. Image processing and fluorescence quantification were all performed using ImageJ software.

### Grafting and root phenotypic analysis

For non-grafted plants, the emerged lateral roots were captured using a Epson dual light scanner v800 with 8-11 dpg seedlings. Lateral root density and primary root length were quantified using ImageJ. For grafting experiments, a transverse cut was made at the upper part of the hypocotyl of 5 dpg seedlings, and combinations between WT and *cyp38-5* scions and root stocks were assembled by attaching the transverse cut surfaces together [38]. These recombined seedlings were then left on wet filter paper plus Hybond N membrane for 5 days in a growth chamber to allow the formation of the junction. After that, the seedlings were transferred back to standard medium for recovery. Scanned root images were taken 7 days after recovery for quantifications of total root branches.

### Protoplast transformation

To better visualize the cellular localization of CYP38 protein, protoplasts were isolated from 3-4 week old Arabidopsis Col-0 plant rosette leaves, and the *proUBQ10:CYP38-Ypet* construct was transformed in using the PEG-calcium method [39]. The transformed protoplasts were then imaged using an SP8 confocal microscope.

### Plant hormones and metabolomics measurements

For plant hormones measurements, 10 days-old WT and *cyp38-5* seedlings grown under standard light conditions and WT grown under low-light conditions were used. For IAA measurements in grafted plants, 20 days-old WT/*cyp38-5* and *cyp38-5*/WT grown under regular light conditions were used. Greater than 100 mg of leaf or root tissue was pulverized into a fine powder with liquid nitrogen and used for LC-MS. Sample loading and data extraction were performed by Creative Proteomics Service (45-1 Ramsey Road, Shirley, NY 11967, USA). For measurements of primary metabolites, 10 days-old WT-0 and *cyp38-5* seedling grown under regular light and WT grown under low light condition were used. Greater than 100 mg of leaf or root tissue were harvested for untargeted GC-MS analysis. Sample loading and data extraction were performed by West Coast Metabolomics Center (UC Davis).

### Westerns blot analysis

Plant tissues were ground under liquid nitrogen to a fine powder. Alkylation assays were performed as previously described [18] using 10 mM methyl-maleimide polyethylene glycol (2-Cys Prx) or 60mM iodoacetamide (FBPase and CYP38). Protein samples were subjected to reducing (2-Cys Prx) or non-reducing (CYP38 and FBPase) SDS-PAGE using acrylamide gel concentration of 9.5% (FBPase and CYP38) and 14% (2-Cys Prx). Resolved proteins were transferred to nitrocellulose membranes and probed with the indicated antibodies. Specific antibody for 2-Cys Prx was previously raised [40]. The anti-FBPase antibody was kindly provided by Dr. Sahrawy (Estación Experimental del Zaidín, Granada, Spain). Antibody for CYP38 was purchased from Agrisera (Sweden).

### Measurements of photosynthetic parameters

Measurement of chlorophyll *a* fluorescence at room temperature was performed using a pulse-amplitude modulation fluorometer (IMAGING-PAM M-Series instrument, Walz, Effeltrich, Germany). Induction–recovery curves were performed using blue (450 nm) actinic light (81 μE m^−2^ s^−1^) at the intensities specified for each experiment during 6 min. Saturating pulses of blue light (10000 μE m^−2^ s^−1^) and 0.6 s duration were applied every 60 s, and recovery in darkness was recorded for another 6 min. The parameters Y(II), Y(NPQ) and Y(NO) corresponding to the respective quantum yields of PSII photochemistry, non-regulated basal quenching and non-regulated energy dissipation, were calculated by the ImagingWin v2.46i software according to the equations described in [41].

## Acknowledgement

We thank the members of Dinneny lab for their discussions during the preparation of this manuscript. Funding was provided by Carnegie Institution for Science Endowment to J.R.D., and European Regional Development Fund-co-financed grant (BIO2017-85195-C2-1-P) from the Spanish Ministry of Economy, Industry and Competitiveness. The plant hormone measurements were provided by Creative Proteomics, and measurements of primary metabolites were supported by West Coast Metabolomics Center in UC Davis. The research of J.R.D. was supported in part by a Faculty Scholar grant from the Howard Hughes Medical Institute and the Simons Foundation.

## Author contributions

L.D., J.M.P., F.J.C. and J.R.D. designed this study. L.D. performed the mutant screen, lateral root phenotypic analysis, cloning, imaging and performed the data analysis. J.M.P. performed the western based studies and measurements of photosynthesis parameters. L.D., J.M.P, F.J.C. and J.R.D. wrote and revised the manuscript.

## Supplementary Information

**Figure S1.**
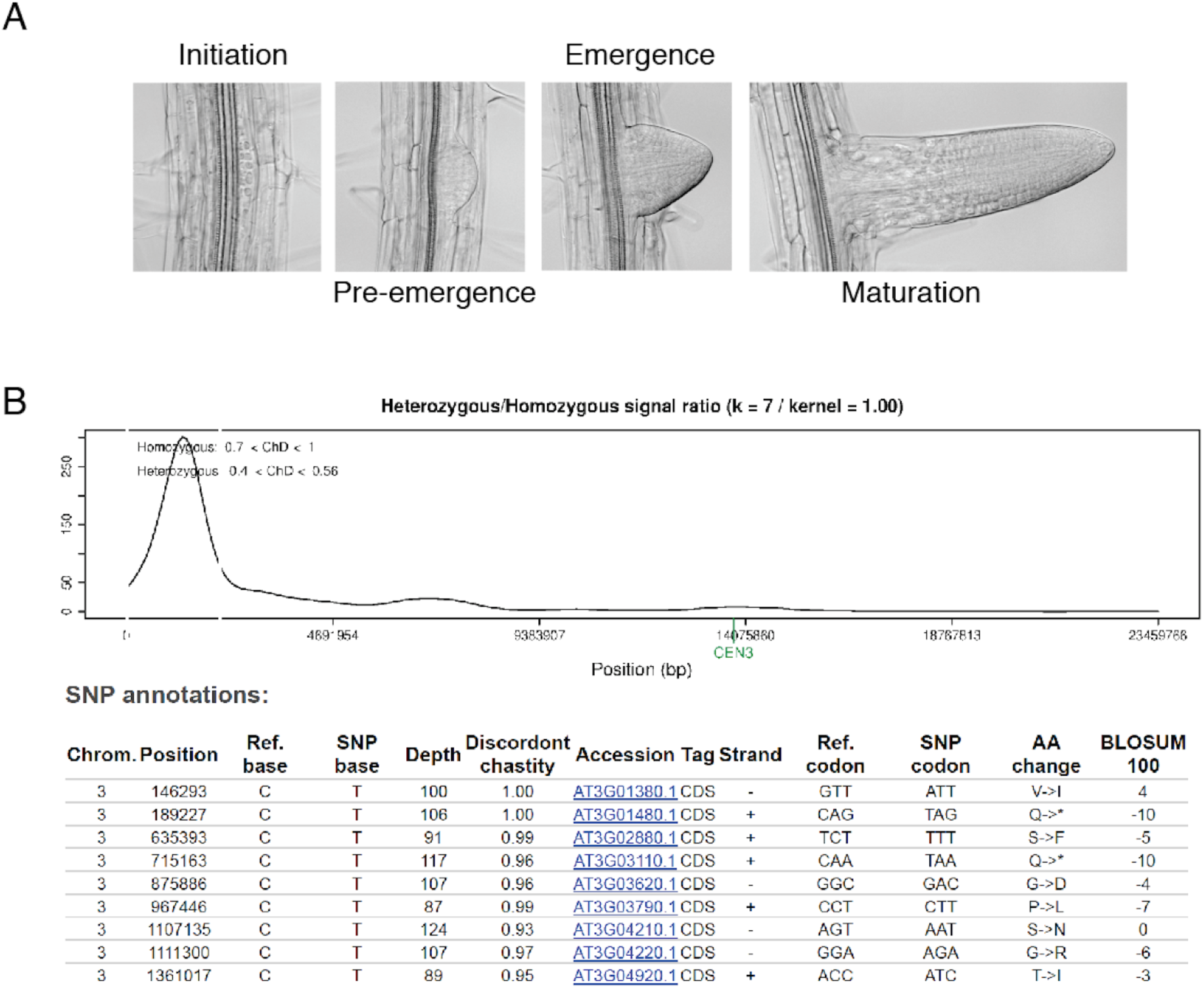
Lateral root developmental stages and mapping results of *cyp38-4*. (A) Representative microscopic images of various stages of lateral roots. (B) NGM (Next-Generation EMS mutation Mapping http://bar.utoronto.ca/ngm/) output of *cyp38-4*. Upper panel is the fine map output showing a peak at the beginning of chromosome 3. Lower panel is the annotations for candidate SNPs. Note that the mutation of *cyp38-4* is the second candidate in the chart.

**Figure S2.**
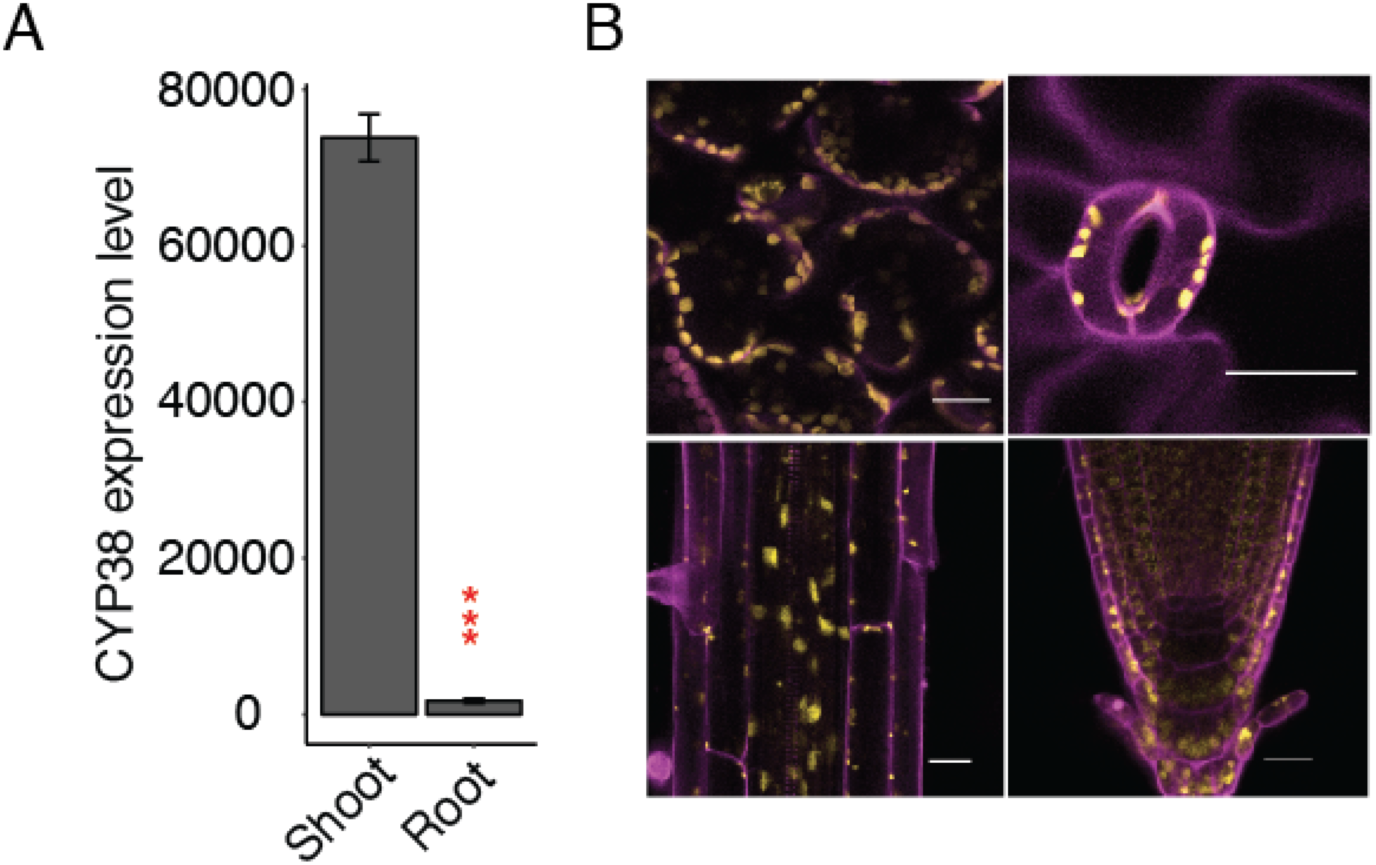
CYP38 expression and localization pattern. (A) Quantitative RT-PCR of *CYP38* transcript level in WT shoot and root tissue. (∗∗∗*P* < 0.001, pairwise T-test with Hochberg correction, n=3) (B) Confocal images of *proUBQ10:CYP38-Ypet* transgene fluorescence in various tissues including mesophyll, stomata, mature root and root tip. Magenta indicates PI staining and autofluorescence from chloroplasts while yellow indicates Ypet fluorescence. Scale bar =20 μm.

**Figure S3.**
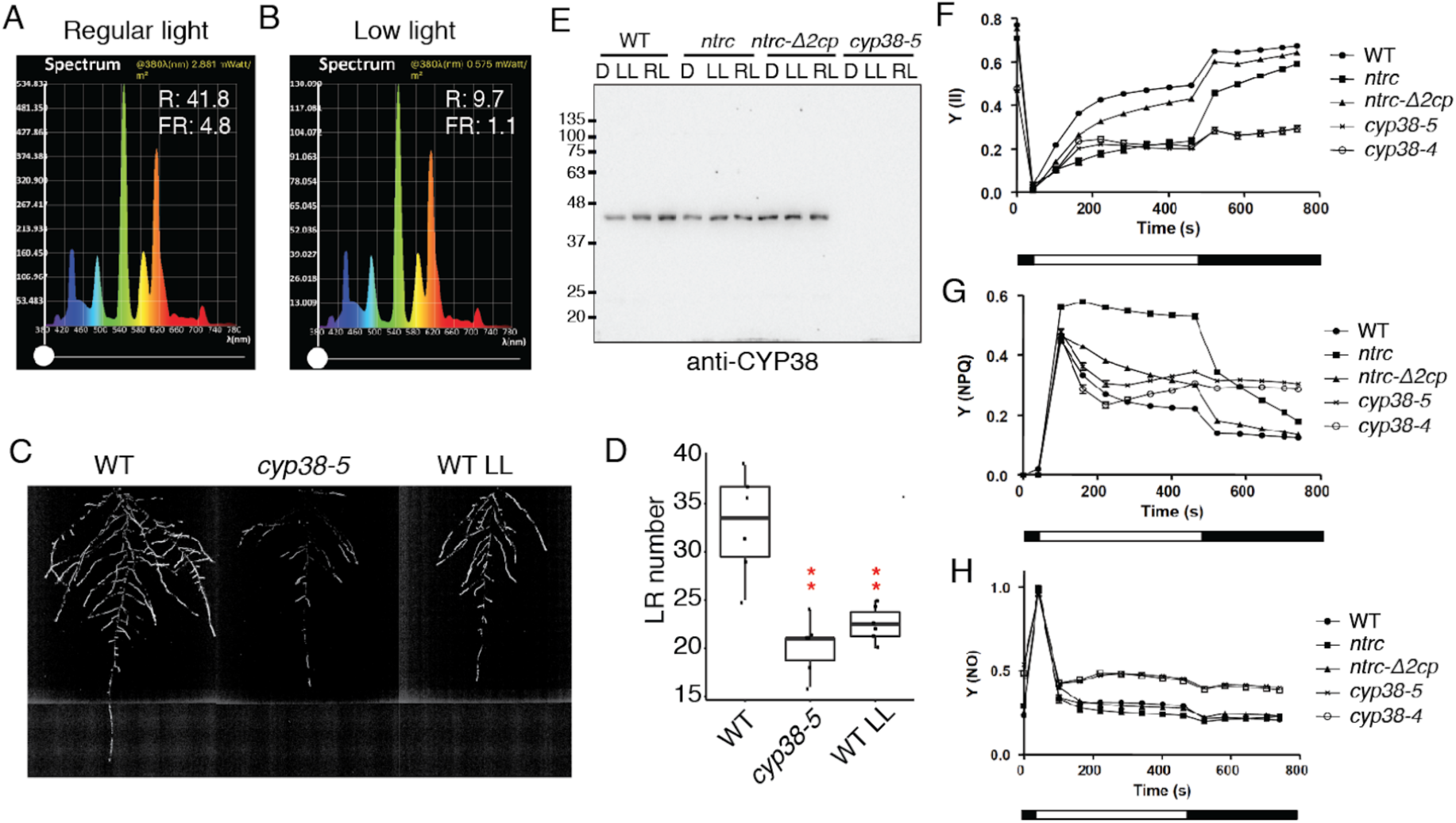
CYP38 dependent root phenotype is derived from photosynthetic status in the shoot. (A, B) Light spectrum measured by Li-Cor spectrometer under regular light (A) and low light (B). R represents Red light level and FR represents Far Red light level. (C, D) Rhizotron images of 21 dpg root system expressing *proUBQ10:Luciferase* (C) and quantifications of lateral root number (D) in WT and *cyp38-5* under regular light and WT under low light. (∗∗*P* < 0.01, pairwise T-test with Hochberg correction, n = 6). (E) Western blot analysis of CYP38 under non-reducing conditions in wild type and mutant plants, as indicated. Protein extracts were obtained from rosette leaves harvested at the end of the dark period (D) and after 30 min of illumination at 50 μE m^−2^ s^−1^ (LL) or 175 μE m^−2^ s^−1^ (RL). Molecular weight markers (kDa) are indicated. (F-H) Measurements of photosynthetic parameters in WT and mutant plants, as indicated. Quantum yields of photosystem II (Y(II)) (F), non-photochemical quenching (Y(NPQ)) (G) and non-regulated energy dissipation ((Y(NO)) (H) were measured in whole plants grown at 125 μE m^−2^ s^−1^ under long-day conditions and adapted to darkness. Determinations were performed four (WT, *ntrc* and *ntrc-*Δ*2cp*) or seven (*cyp38-4* and *cyp38-5*) times and each data point is the mean ± SE. White and black blocks indicate periods of illumination with actinic light (81 mE m^−2^ s^−1^) and darkness, respectively.

**Figure S4.**
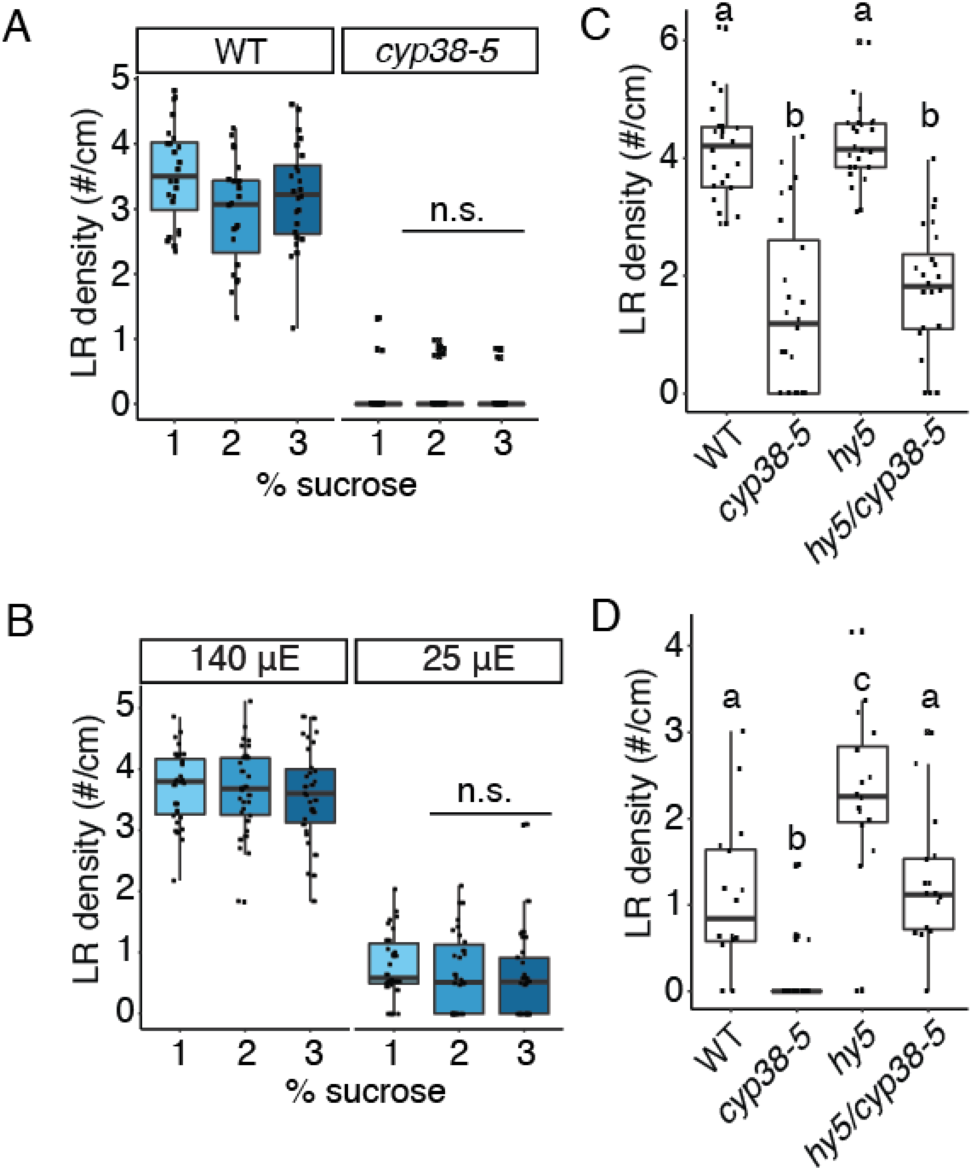
Sucrose and hy5 signaling are not necessary for CYP38 dependent lateral root emergence. (A, B) Quantifications of lateral root density comparing WT and *cyp38-5* under different levels of sucrose in the growth media (A), or comparing WT under regular and low light conditions and with different levels of sucrose in the growth media (B). (n.s. represents not significant, two-way ANOVA comparing with WT under different sucrose concentrations, n>23) (C, D) Quantifications of lateral root density comparing WT, *cyp38-5, hy5 and hy5/cyp38-5* under regular light, 140 μE m^−2^ s^−1^ (A), and low light, 25 μE m^−2^ s^−1^ (B). Significantly different groups are indicated with letters (*P* < 0.05), n>16.

**Figure S5.**
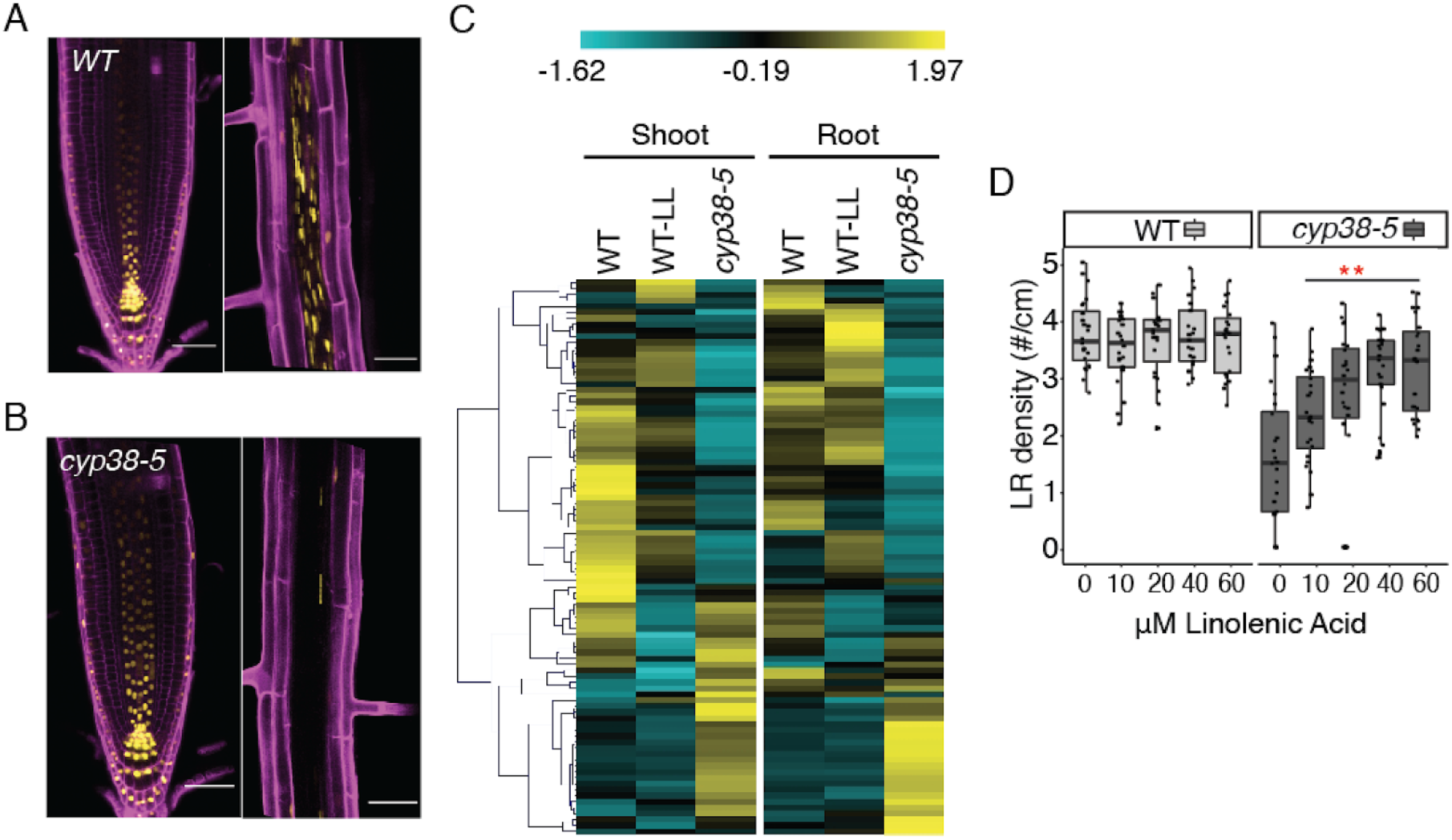
Auxin and defense related metabolites are affected in *cyp38* mutant. (A, B) Confocal images of *pDR5rev::3XVenus-N7* in primary root tip (left panel) and mature region of primary root (right panel) in WT (A), or in *cyp38-5* (B). Scale bar =50 μm. (C)Heatmap indicating the levels of various primary metabolites in WT, WT under low light, and *cyp38-5*, in both shoot and root. Hierarchical clustering was used to group metabolites with similar trends. Values are taken from the average of 3 individual measurements. (D) Quantification of lateral root density in WT compared with *cyp38-5* with exogenous linolenic acid treatment. (∗∗*P* < 0.01, two-way ANOVA comparing WT under standard conditions and linolenic acid treatment. n>21)

**Table S1.**
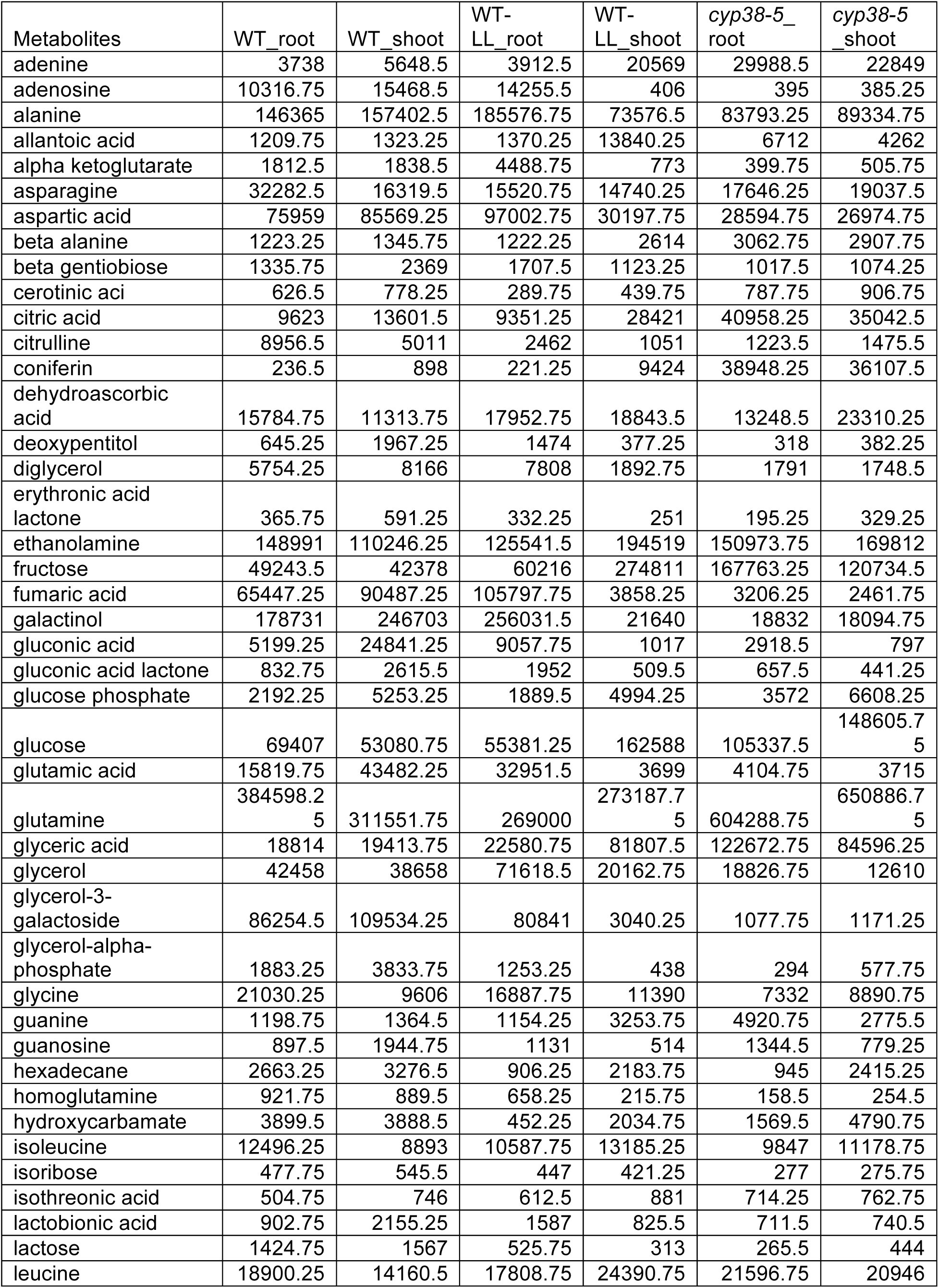
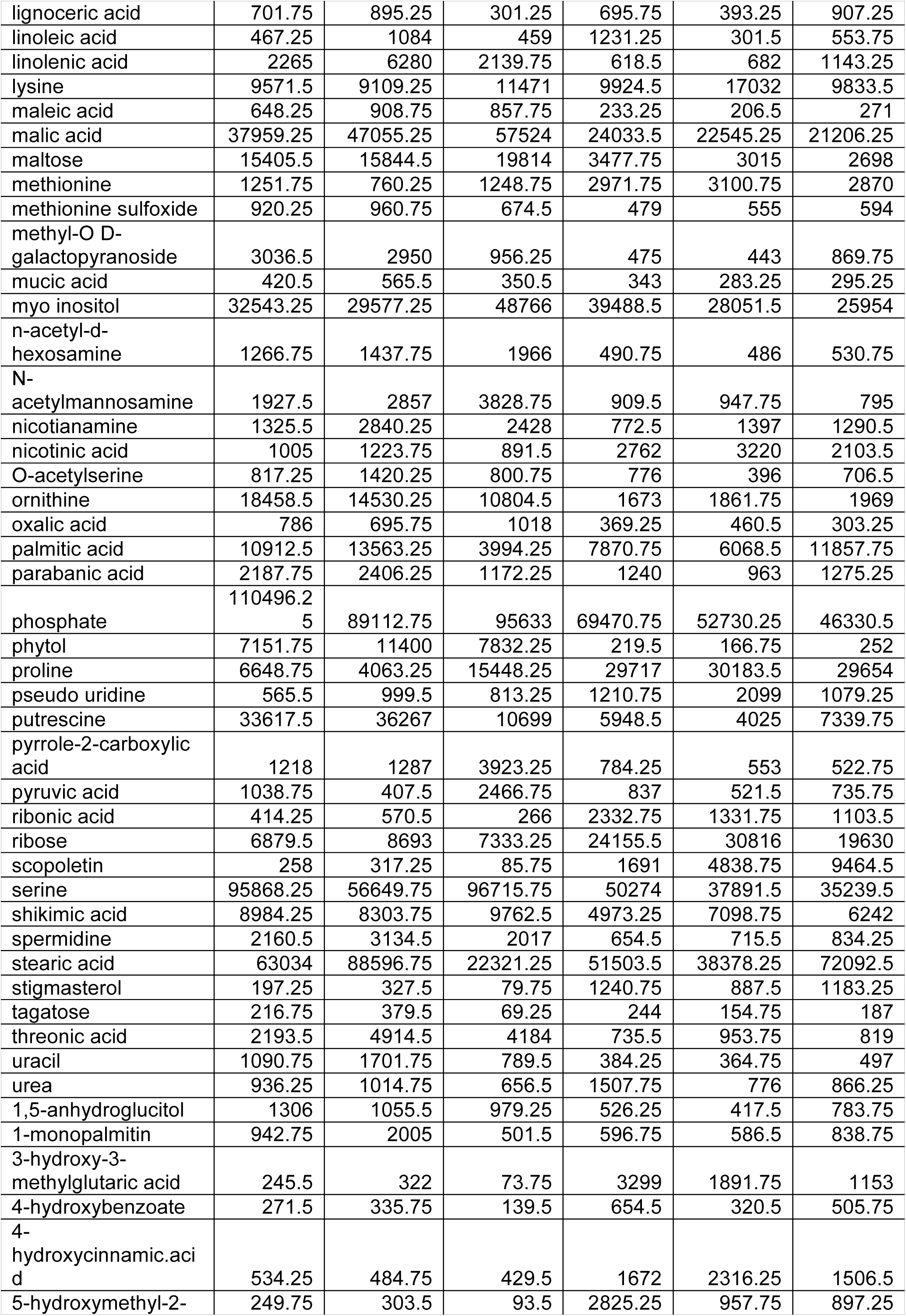
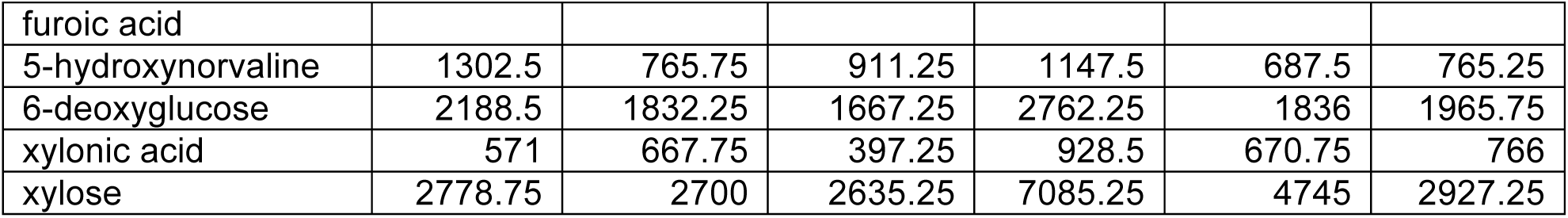
Levels of various primary metabolites in WT, WT under low light, and *cyp38-5*.

**Table S2.** Metabolites that are significantly lower in both WT-LL and *cyp38-5*.

| Metabolites | WT_shoot | WT-LL_shoot | *cyp38-5*_shoot | WT_root | WT-LL_root | *cyp38-5*_root |
| --- | --- | --- | --- | --- | --- | --- |
| galactinol | 246703 | 186135.25 | 18094.75 | 178731 | 256031.5 | 18832 |
| gluconic acid | 24841.25 | 8821 | 797 | 5199.25 | 9057.75 | 2918.5 |
| glycerol-3-galactoside | 109534.25 | 81119.75 | 1171.25 | 86254.5 | 80841 | 1077.75 |
| glycerol-alpha-phosphate | 3833.75 | 1409.5 | 577.75 | 1883.25 | 1253.25 | 294 |
| linolenic acid | 6280 | 1334 | 1143.25 | 2265 | 2139.75 | 682 |
| parabanic acid | 2406.25 | 1139.5 | 1275.25 | 2187.75 | 1172.25 | 963 |
| phytol | 11400 | 6798 | 252 | 7151.75 | 7832.25 | 166.75 |
| spermidine | 3134.5 | 1371 | 834.25 | 2160.5 | 2017 | 715.5 |
| tagatose | 379.5 | 85 | 187 | 216.75 | 69.25 | 154.75 |
| uracil | 1701.75 | 639.5 | 497 | 1090.75 | 789.5 | 364.75 |
| agmatine | 505.5 | 262.5 | 336.25 | 411 | 118 | 167.5 |
| putrescine | 36267 | 21212.25 | 7339.75 | 33617.5 | 10699 | 4025 |

**Table S3.**
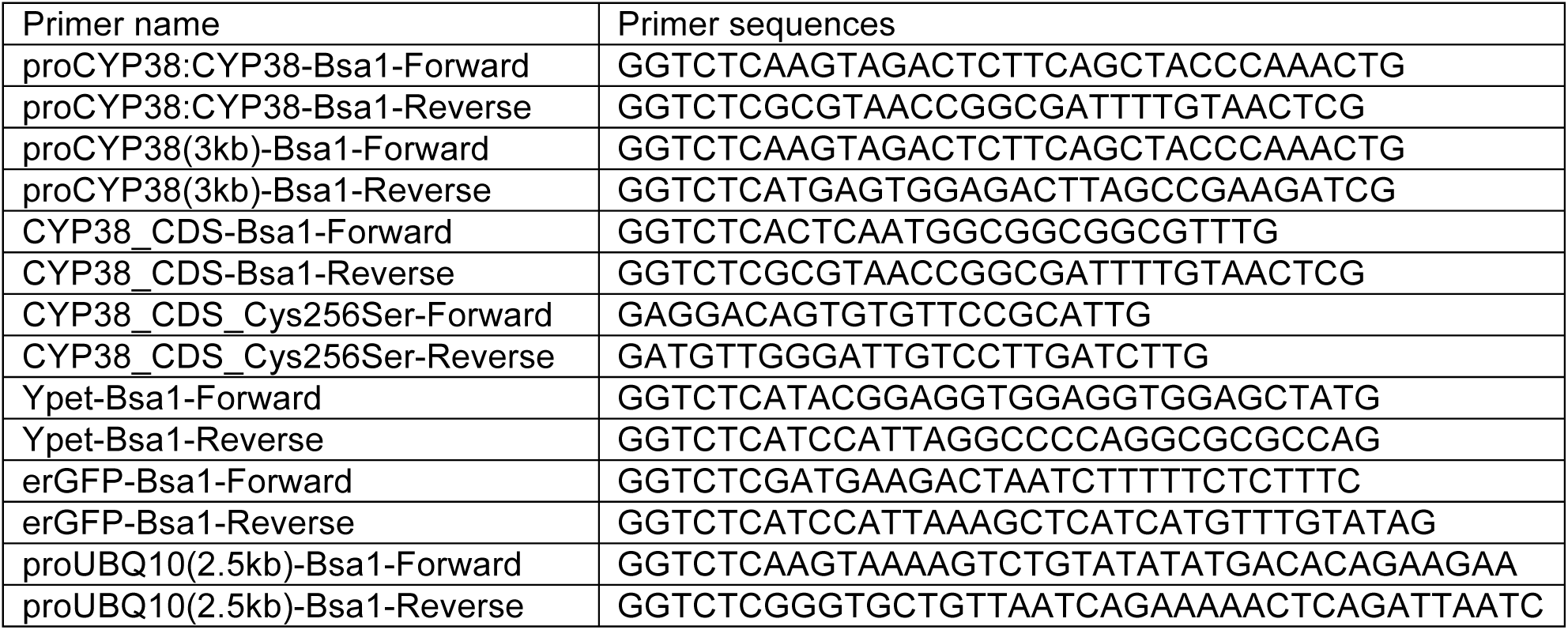
Primers for cloning.

## References

1. Knoblauch, M., and Peters, W.S. (2013). Long-distance translocation of photosynthates: a primer. Photosynth. Res. 117, 189–196.

2. Kircher, S., and Schopfer, P. (2012). Photosynthetic sucrose acts as cotyledon-derived long-distance signal to control root growth during early seedling development in Arabidopsis. Proc. Natl. Acad. Sci. U. S. A. 109, 11217–11221.

3. Fu, A., He, Z., Cho, H.S., Lima, A., Buchanan, B.B., and Luan, S. (2007). A chloroplast cyclophilin functions in the assembly and maintenance of photosystem II in Arabidopsis thaliana. Proc. Natl. Acad. Sci. U. S. A. 104, 15947–15952.

4. Mashiguchi, K., Tanaka, K., Sakai, T., Sugawara, S., Kawaide, H., Natsume, M., Hanada, A., Yaeno, T., Shirasu, K., Yao, H., et al. (2011). The main auxin biosynthesis pathway in Arabidopsis. Proc. Natl. Acad. Sci. U. S. A. 108, 18512–18517.

5. Zhao, Y. (2012). Auxin biosynthesis: a simple two-step pathway converts tryptophan to indole-3-acetic acid in plants. Mol. Plant 5, 334–338.

6. Van Norman, J.M., Xuan, W., Beeckman, T., and Benfey, P.N. (2013). To branch or not to branch: the role of pre-patterning in lateral root formation. Development 140, 4301–4310.

7. Swarup, K., Benkova, E., Swarup, R., Casimiro, I., Peret, B., Yang, Y., Parry, G., Nielsen, E., De Smet, I., Vanneste, S., et al. (2008). The auxin influx carrier LAX3 promotes lateral root emergence. Nat. Cell Biol. 10, 946–954.

8. Péret, B., Larrieu, A., and Bennett, M.J. (2009). Lateral root emergence: a difficult birth. J. Exp. Bot. 60, 3637–3643.

9. Fernández-Marcos, M., Desvoyes, B., Manzano, C., Liberman, L.M., Benfey, P.N., Del Pozo, J.C., and Gutierrez, C. (2017). Control of Arabidopsis lateral root primordium boundaries by MYB36. New Phytol. 213, 105–112.

10. Vasudevan, D., Fu, A., Luan, S., and Swaminathan, K. (2012). Crystal structure of Arabidopsis cyclophilin38 reveals a previously uncharacterized immunophilin fold and a possible autoinhibitory mechanism. Plant Cell 24, 2666–2674.

11. SirpiÃ¶, S., Khrouchtchova, A., Allahverdiyeva, Y., Hansson, M., Fristedt, R., Vener, A.V., Scheller, H.V., Jensen, P.E., Haldrup, A., and Aro, E.-M. (2008). AtCYP38 ensures early biogenesis, correct assembly and sustenance of photosystem II. Plant J. 55, 639–651.

12. Ivanchenko, M.G., Zhu, J., Wang, B., Medvecká, E., Du, Y., Azzarello, E., Mancuso, S., Megraw, M., Filichkin, S., Dubrovsky, J.G., et al. (2015). The cyclophilin A DIAGEOTROPICA gene affects auxin transport in both root and shoot to control lateral root formation. Development 142, 712–721.

13. Spiegelman, Z., Ham, B.-K., Zhang, Z., Toal, T.W., Brady, S.M., Zheng, Y., Fei, Z., Lucas, W.J., and Wolf, S. (2015). A tomato phloem-mobile protein regulates the shoot-to-root ratio by mediating the auxin response in distant organs. Plant J. 83, 853–863.

14. Duan, L., Dietrich, D., Ng, C.H., Chan, P.M.Y., Bhalerao, R., Bennett, M.J., and Dinneny, J.R. (2013). Endodermal ABA signaling promotes lateral root quiescence during salt stress in Arabidopsis seedlings. Plant Cell 25, 324–341.

15. Rellán-Álvarez, R., Lobet, G., Lindner, H., Pradier, P.-L., Sebastian, J., Yee, M.-C., Geng, Y., Trontin, C., LaRue, T., Schrager-Lavelle, A., et al. (2015). GLO-Roots: an imaging platform enabling multidimensional characterization of soil-grown root systems. Elife 4. Available at: http://dx.doi.org/10.7554/eLife.07597.

16. Kirchsteiger, K., Ferrández, J., Pascual, M.B., González, M., and Cejudo, F.J. (2012). NADPH thioredoxin reductase C is localized in plastids of photosynthetic and nonphotosynthetic tissues and is involved in lateral root formation in Arabidopsis. Plant Cell 24, 1534–1548.

17. Serrato, A.J., Pérez-Ruiz, J.M., Spínola, M.C., and Cejudo, F.J. (2004). A novel NADPH thioredoxin reductase, localized in the chloroplast, which deficiency causes hypersensitivity to abiotic stress in Arabidopsis thaliana. J. Biol. Chem. 279, 43821–43827.

18. Pérez-Ruiz, J.M., Naranjo, B., Ojeda, V., Guinea, M., and Cejudo, F.J. (2017). NTRC-dependent redox balance of 2-Cys peroxiredoxins is needed for optimal function of the photosynthetic apparatus. Proc. Natl. Acad. Sci. U. S. A. 114, 12069–12074.

19. Cerveau, D., Kraut, A., Stotz, H.U., Mueller, M.J., Couté, Y., and Rey, P. (2016). Characterization of the Arabidopsis thaliana 2-Cys peroxiredoxin interactome. Plant Sci. 252, 30–41.

20. Wang, Y., Zeng, L., and Xing, D. (2015). ROS-mediated enhanced transcription of CYP38 promotes the plant tolerance to high light stress by suppressing GTPase activation of PsbO2. Front. Plant Sci. 6, 777.

21. Macgregor, D.R., Deak, K.I., Ingram, P.A., and Malamy, J.E. (2008). Root system architecture in Arabidopsis grown in culture is regulated by sucrose uptake in the aerial tissues. Plant Cell 20, 2643–2660.

22. Chen, X., Yao, Q., Gao, X., Jiang, C., Harberd, N.P., and Fu, X. (2016). Shoot-to-Root Mobile Transcription Factor HY5 Coordinates Plant Carbon and Nitrogen Acquisition. Curr. Biol. 26, 640–646.

23. Zhang, Y., Wang, C., Xu, H., Shi, X., Zhen, W., Hu, Z., Huang, J., Zheng, Y., Huang, P., Zhang, K.-X., et al. (2019). HY5 Contributes to Light-Regulated Root System Architecture Under a Root-Covered Culture System. Front. Plant Sci. 10, 1490.

24. van Gelderen, K., Kang, C., Paalman, R., Keuskamp, D., Hayes, S., and Pierik, R. (2018). Far-Red Light Detection in the Shoot Regulates Lateral Root Development through the HY5 Transcription Factor. Plant Cell 30, 101–116.

25. Du, Y., and Scheres, B. (2018). Lateral root formation and the multiple roles of auxin. J. Exp. Bot. 69, 155–167.

26. Lavenus, J., Goh, T., Roberts, I., Guyomarc’h, S., Lucas, M., De Smet, I., Fukaki, H., Beeckman, T., Bennett, M., and Laplaze, L. (2013). Lateral root development in Arabidopsis: fifty shades of auxin. Trends Plant Sci. 18, 450–458.

27. Heisler, M.G., Ohno, C., Das, P., Sieber, P., Reddy, G.V., Long, J.A., and Meyerowitz, E.M. (2005). Patterns of auxin transport and gene expression during primordium development revealed by live imaging of the Arabidopsis inflorescence meristem. Curr. Biol. 15, 1899–1911.

28. Cha, J.-Y., Kim, W.-Y., Kang, S.B., Kim, J.I., Baek, D., Jung, I.J., Kim, M.R., Li, N., Kim, H.-J., Nakajima, M., et al. (2015). A novel thiol-reductase activity of Arabidopsis YUC6 confers drought tolerance independently of auxin biosynthesis. Nat. Commun. 6, 8041.

29. Geigenberger, P., and Fernie, A.R. (2014). Metabolic control of redox and redox control of metabolism in plants. Antioxid. Redox Signal. 21, 1389–1421.

30. Cejudo, F.J., Ojeda, V., Delgado-Requerey, V., González, M., and Pérez-Ruiz, J.M. (2019). Chloroplast Redox Regulatory Mechanisms in Plant Adaptation to Light and Darkness. Front. Plant Sci. 10, 380.

31. de Werra, P., Péchy-Tarr, M., Keel, C., and Maurhofer, M. (2009). Role of gluconic acid production in the regulation of biocontrol traits of Pseudomonas fluorescens CHA0. Appl. Environ. Microbiol. 75, 4162–4174.

32. Chanda, B., Xia, Y., Mandal, M.K., Yu, K., Sekine, K.-T., Gao, Q.-M., Selote, D., Hu, Y., Stromberg, A., Navarre, D., et al. (2011). Glycerol-3-phosphate is a critical mobile inducer of systemic immunity in plants. Nat. Genet. 43, 421–427.

33. Seifi, H.S., and Shelp, B.J. (2019). Spermine Differentially Refines Plant Defense Responses Against Biotic and Abiotic Stresses. Front. Plant Sci. 10, 117.

34. Lou, Y.-R., Bor, M., Yan, J., Preuss, A.S., and Jander, G. (2016). Arabidopsis NATA1 Acetylates Putrescine and Decreases Defense-Related Hydrogen Peroxide Accumulation. Plant Physiol. 171, 1443–1455.

35. Vellosillo, T., Martínez, M., López, M.A., Vicente, J., Cascón, T., Dolan, L., Hamberg, M., and Castresana, C. (2007). Oxylipins produced by the 9-lipoxygenase pathway in Arabidopsis regulate lateral root development and defense responses through a specific signaling cascade. Plant Cell 19, 831–846.

36. Emami, S., Yee, M.-C., and Dinneny, J.R. (2013). A robust family of Golden Gate Agrobacterium vectors for plant synthetic biology. Front. Plant Sci. 4, 339.

37. Song, L., Huang, S.-S.C., Wise, A., Castanon, R., Nery, J.R., Chen, H., Watanabe, M., Thomas, J., Bar-Joseph, Z., and Ecker, J.R. (2016). A transcription factor hierarchy defines an environmental stress response network. Science 354. Available at: http://dx.doi.org/10.1126/science.aag1550.

38. Melnyk, C.W. (2017). Grafting with Arabidopsis thaliana. Methods Mol. Biol. 1497, 9–18.

39. Yoo, S.-D., Cho, Y.-H., and Sheen, J. (2007). Arabidopsis mesophyll protoplasts: a versatile cell system for transient gene expression analysis. Nature Protocols 2, 1565–1572. Available at: http://dx.doi.org/10.1038/nprot.2007.199.

40. Pérez-Ruiz, J.M., Spínola, M.C., Kirchsteiger, K., Moreno, J., Sahrawy, M., and Cejudo, F.J. (2006). Rice NTRC is a high-efficiency redox system for chloroplast protection against oxidative damage. Plant Cell 18, 2356–2368.

41. Kramer, D.M., Johnson, G., Kiirats, O., and Edwards, G.E. (2004). New Fluorescence Parameters for the Determination of QA Redox State and Excitation Energy Fluxes. Photosynth. Res. 79, 209.

